# Process-based model predicts seasonal variation in eDNA transport - a case study on Eurasian beavers in a small river

**DOI:** 10.64898/2026.02.04.703803

**Authors:** James A. Macarthur, Didier Pont, Meriem Bouhouche, Barbara Morrisey, Nathan P. Griffiths, Frank Rosell, Jørn Henrik Sønstebø, Martin J. Gaywood, Bernd Hänfling

## Abstract

Robust methods to monitor species distributions are vital to ensuring successful conservation strategies, particularly in the case of conservation translocations. Environmental DNA (eDNA) from water samples is a cost-effective method to monitor species distributions without physical capture or disturbance. However, eDNA is vulnerable to long-distance transport depending on the hydrological and environmental characteristics which can lead to spatially false positives and ultimately inaccurate species distributions. Recently, the development of particle transport models has allowed researchers to integrate hydrological and environmental variables to predict how far eDNA will transport from a source point. Here, eDNA samples (n=218) were collected and quantified using digital PCR (dPCR) to study monthly changes in Eurasian beaver (*Castor fiber*) eDNA concentrations downstream of an enclosure which contained 4 - 5 beavers located in Scotland. The shortest eDNA transport distances (< 2 km) were observed in the summer which correlated with the lowest flows and highest temperatures. In contrast, throughout the winter eDNA was consistently detected up to 5.8 km downstream correlating with the highest discharge and lowest temperature. The eDNA transport model reliably reproduced the decrease in eDNA concentrations downstream of the enclosure, however there were challenges surrounding stream-specific decay rates following a confluence. To study localised species distributions, samples should be collected during summer low flow conditions. Conversely, to maximise species detections sampling should be conducted in winter which had the longest eDNA transport and highest detectability.

## Introduction

Environmental DNA (eDNA) based approaches have become a popular tool to monitor species distributions where its high-sensitivity and ability to detect species without physical capture or disturbance provides a distinct advantage to traditional monitoring techniques (Burgher *et al*., 2024; Kalogianni et al., 2024; Wood et al., 2025). Molecular methods are particularly effective when monitoring rare (Griffiths *et al*., 2020, 2023; Kalogianni et al., 2024), invasive (Cook *et* al., 2025; Curtis *et al*., 2021) or translocated species (Burgher et al., 2024; Wood et al., 2025). These approaches are subject to limitations, particularly in lotic systems, where downstream transport of eDNA away from its source complicates interpretation of the spatial scale and can lead to false positives (Deiner & Altermatt, 2014; Goldberg et al., 2016; Pont *et al*., 2018; Wacker et al., 2019). Experiments using caged brown trout (*Salmo trutta*) revealed stream-specific variation in eDNA transport, where higher flows had a negative effect on eDNA copy numbers in a larger stream with diverse flows; conversely, in a smaller stream higher flows resulted in minimal decrease in eDNA copies with distance (Jane *et al*., 2015). Subsequently, stream mesocosm experiments demonstrated how stream substrate can impact eDNA retention and subsequent transport (Shogren et al., 2017). Rapid transport and no reduction in eDNA concentrations was observed in cobble sections, conversely the shortest transport was observed in fine sediment sections related to both sedimentation, resuspension and absorption processes (Shogren *et al*., 2017). Finally, eDNA transportation distances for fish ranged from a few km in smaller streams up to over 100 km in large river systems (Pont *et al*., 2018). Irrespective of this long distance eDNA transport, consistent spatial patterns of the fish community could be drawn between eDNA and traditional electrofishing surveys (Pont et al., 2018).

Seasonal variation in eDNA concentrations and the subsequent transport has been shown to be dependent on species behaviour, ecology, DNA shedding rates and environmental conditions (Griffiths et al., 2025; Jo & Yamanaka, 2022; Reves et al., 2026). Reproductive events are often associated with a marked increase in eDNA concentration (Di Muri *et al*., 2023; Wacker et al., 2019). The release of larvae between May and August caused a 20-fold increase in freshwater pearl mussel (*Margaritifera margaritifera*) eDNA concentrations, however irrespective of this seasonal increase, eDNA concentrations remained consistently high downstream throughout the subsequent 1.7 km of river (Wacker et al., 2019). Similarly, amphibians from streams in Idaho were harder to detect in Spring than Autumn, which could be a result of reduced metabolism in colder weather, changes in species behaviour or seasonal changes in species density (Goldberg et al., 2011). Furthermore, several studies have demonstrated a seasonal increase in terrestrial vertebrate detections and eDNA transport during the rainy season (Bai et al., 2025; Condachou et al., 2025). Recently, the development of particle transport models have allowed researchers to integrate hydrological and environmental variables to predict how far eDNA will transport from a source point (Carraro et al., 2023; Pont, 2024; URycki et al., 2024). A 30-fold increase in spatial resolution was highlighted when integrating eDNA transport models to characterise catchment-scale biodiversity patterns for fish, invertebrates and bacterial communities (Carraro *et al*., 2023). Subsequent research demonstrated the utility of these eDNA transport models to upscale ecological indices for macroinvertebrates (Blackman et al., 2024). Finally, a process-based downstream transport model was developed to predict fish eDNA transport distances as a function of hydraulic conditions, eDNA transfer to the bottom and decay rate of target eDNA molecules (Pont, 2024). In the Danube River, predicted and observed eDNA concentrations of grayling (*Thymallus thymallus*) were consistent 82 km downstream of the confluence with the Lech tributary, despite electrofishing confirming grayling were absent from this section of the main river (Pont, 2024).

Conservation translocations are now an increasingly accepted methodology to reinforce extirpated or threatened species and restore ecosystem functions (Gaywood, 2024). A recent review cited the most common challenge following translocations was the movement and dispersal of individuals, notably long-distance dispersal from the intended release sites which could reduce the risk of success (Berger-Tal et al., 2020). Continuous monitoring post-release is vital to ensure successful translocations (Bubac et al., 2019; Gaywood, 2024; Stone et al., 2025). However, this can be labour intensive, expensive and intrusive particularly when dealing with elusive species, remote release sites and long-distance dispersal (Berger-Tal *et* al., 2020; Bilby & Moseby, 2024). Beavers (*Castor spp*.) as a keystone species have become a popular target for conservation translocation to promote ecological functioning and restore aquatic biodiversity (Brazier et al., 2021; Fairfax & Westbrook, 2024; Law et al., 2017; Rosell et al., 2005). Traditional monitoring post beaver translocation has included physical recapture of released individuals to take genetic samples (Taylor et al., 2024) or to screen individual health (Campbell-Palmer et al., 2021a), telemetry surveys to study individual dispersal and behavioural patterns (Doden et al., 2022), camera traps (Dytkowicz *et al*., 2023) and visual observation (Rosell & Nolet, 1997) to identify individuals, aerial and satellite imagery to study impacts of beaver modifications on the wider ecosystem (Fairfax et al., 2023; Puttock *et al*., 2015) and finally, field sign surveys to study foraging preferences, identify habitat modifications, characterise territories and estimate population sizes (Campbell-Palmer et al., 2021b; Mori et al., 2024). Field sign surveys have been the primary monitoring method in Scotland (Campbell-Palmer et al., 2021b, 2021c; Campbell *et al*., 2012); however these surveys become logistically challenging as beaver populations grow, particularly due to their territorial behaviour and occasional large dispersal distances (> 20 km) (Mayer *et al*., 2017a; Sun et al., 2000).

In recent years, studies have demonstrated that eDNA from water samples could reliably detect both Eurasian (*C. fiber*) and North American beavers (*C. canadensis*) in captive (Broadhurst et al., 2021; Harper et al., 2019), wild (Bobeva et al., 2025; Macarthur et al., 2025) and translocated populations (Burgher *et al*., 2024; Duke *et al*., 2025). Following translocations in Washington State, North American beaver eDNA could be rapidly detected up to 2.9 km from the release sites within 1 week, highlighting the potential for long distance eDNA transport (Burgher et al., 2024). Subsequent surveys demonstrated that beaver eDNA could be detected even without visual field signs (Duke et al., 2025). Finally, research from Scotland, demonstrated consistently high detections of 79% across three replicates for the Eurasian beaver in Scotland using eDNA (Macarthur et al., 2025). Further comparisons between positive beaver eDNA and historic beaver field signs (Campbell-Palmer et al., 2021a) revealed an 82.4 % overlap, whilst this survey also identified a large number of areas where beavers have expanded since the previous survey (Macarthur *et al*., 2025).

Previously eDNA transport models have primarily been applied to fish (Pont, 2024) and aquatic invertebrate species (Blackman et al., 2024; Carraro et al., 2023), meanwhile the relationship between streamflow and mammal eDNA transport remains understudied (Bai et al., 2025). The consistent detections of beavers from previous eDNA surveys makes them an ideal candidate for validating eDNA transport models on semi-aquatic mammals (Broadhurst *et al*., 2021; Burgher et al., 2024; Harper et al., 2019; Macarthur et al., 2025). The aim of the study was to understand seasonal variation of beaver eDNA transport in a small stream and to test if it can be predicted using a process-based transport model. We took advantage of a beaver enclosure which provided us with a spatially explicit source of beaver eDNA feeding into a natural stream. We quantified beaver eDNA concentration across the catchment over the course of a year and integrated an eDNA transport model to predict downstream transport relative to environmental and hydrological characteristics. We hypothesized that eDNA transport would be highly correlated with discharge and temperature, therefore we predict the longest eDNA transport in winter and the shortest transport in summer.

## Methods

### Study region

This study was carried out within the Moniack catchment in the Scottish Highlands where a 40-hectare enclosure has supported a small population of 4 - 5 Eurasian beavers since 2008 (Figure 1). Immediately downstream of the enclosure flows a small river (hereafter small river) which runs for approximately 1.5 km with an average width of 2.3 metres and an average depth of 0.13 m before entering the larger Moniack river (hereafter Moniack river) which runs for a further 5.6 km with an average width of 5.1 metres and an average depth of 0.16 m before entering the Beauly Firth. Sites ranged in elevation from 155 m to 4 m and the water temperatures throughout the sampling period ranged from 19.1 °C in July to 2.3 °C in December. Agriculture is the primary land-use throughout the surrounding area, however sites 5, 6 and 7 are situated within a mixed woodland forest. See Table S1 for full details on site locations.

**Figure 1:**
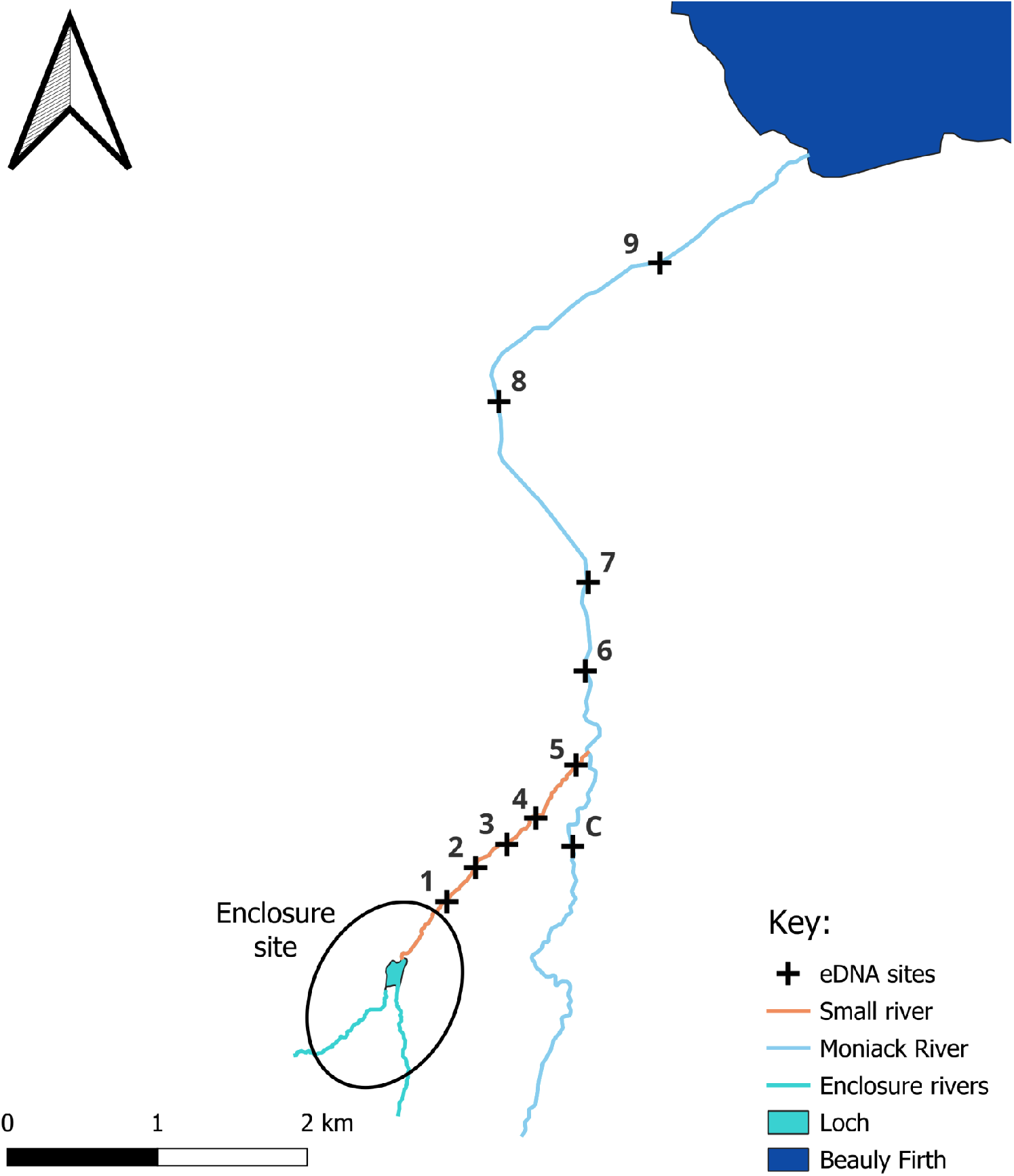
eDNA water sampling locations (n = 10, black cross) downstream of an enclosure site ranging from 0 to 5.8 km. Two water samples were collected from a control site (C) upstream of the confluence between the small river and the Moniack river to confirm there were no other sources of beaver eDNA within the wider catchment.

### Sample collection and filtration

Between 06/08/2024 - 03/07/2025, 216 2-litre surface samples were collected from nine sites (two field replicates per site) up to 5.8 km downstream of the beaver enclosure (Fig. 1). No wild beaver population is known in this catchment. Therefore, the only source of beaver eDNA is from the enclosure site. An additional two water samples were collected from a control site (C) in August 2024 upstream of the confluence between the small river and the Moniack river to confirm there were no other sources of beaver eDNA within the wider catchment. Each site was visited monthly over the course of a year, and months were assigned to the following seasons: Summer (June, July, August), Autumn (September, October, November), Winter (December, January, February) and Spring (March, April, May). Sample collection and filtration followed the methodology described in (Griffiths et al., 2023). Field blanks (n = 13) of purified water were used each visit to monitor contamination. Water samples were vacuum-filtered within 24 hours of collection through sterile 0.45 μm mixed cellulose nitrate membrane filters with pads (47 mm diameter; Whatman, GE Healthcare) using Pall filtration units prior to being stored at -20 °C. See methods S1 for full details. Ethical approval for the current study was obtained under the application “ETH2223-0838” to UHI.

### Collection of environmental covariates

Stream discharge was measured each month at four sites (1, 5, 6 and 9) using the OTT MF Pro flow meter (OTT HydroMet Corporation, Virginia, United States). Briefly, the stream velocity, depth and distance across a transect was measured at regular intervals across the stream cross-section, and the number of stations was adjusted accordingly to stream width following the manufacturer guidelines. Water temperature was recorded at each site using the HI-98129 probe (Hanna Instruments LTD, Bedfordshire, UK) (Table S2).

### DNA extractions

DNA was extracted using the modified mu-DNA lysis water protocol (Sellers *et al*., 2018) described in Macarthur *et al*., (2025). Extraction blanks (n = 12) consisting of empty tubes and extraction reagents were used each month to monitor contamination.

### Digital PCR

Digital PCR was performed in duplicate (2 x PCR replicates per eDNA extract) using the following unpublished primers F (5’-CCTCGGTGCCATCAACTTTA-3’) and R (5’-GCAGTAACTAGGACGGATCATAC-3’) to amplify a 102 bp fragment of the Eurasian beaver COX1 gene (Bouhouche, 2023). These primers were previously validated *in vitro, in silico* and *in situ* to ensure high specificity to the Eurasian beaver using qPCR (Bouhouche, 2023). A positive control (quantified at 0.01 ng/μl) of Eurasian beaver tissue and a negative control of molecular grade water were used for each plate and the TaqMan Exogenous Internal Positive Control (IPC) was used to test for inhibition (Applied Biosystems, Warrington, UK). Reaction volumes consisted of 13.3 μl QIAcuity EG PCR kit (QIAGEN, Manchester, UK) 1 μl primers (0.5 x each 10 μM Primer (Integrated DNA Technologies, Belgium), 4 μl IPC Mix and 0.4 μl IPC DNA (Applied Biosystems, Warrington, UK), 16.3 μl of molecular grade water and 5 μl of DNA extract.

Reactions were run on the QIAcuity-00896 digital PCR system (QIAGEN, Manchester, UK) using the 26k nanowell plates with the following thermocycling profile: 95°C for 10 min, 35 cycles of 95°C for 15 s and 60°C for 1 min. The beaver and IPC assays used the EVAGreen dye and VIC dye respectively and imaging for both channels was conducted at an exposure duration of 300 ms and gain 6. As the EVAGreen assay fluoresces both beaver eDNA and the IPC a common threshold was set at 130 RFU for the beaver channel to ensure only beaver eDNA was retained (Figure S1), meanwhile the auto-threshold was used to quantify the IPC. The limit of detection (LOD) was calculated following a discrete threshold approach which ran three different concentrations of beaver tissue aiming to achieve ≧95% detections across 20 replicates (Klymus et al., 2020).

### Data analysis

All maps were created in QGIS (QGIS Development Team, 2025), analyses and graphical visualisation used R v.4.5.2 (R Core Team, 2025) with ggplot2(Wickham, 2016). Initially, a Spearman’s rank correlation was conducted to test the relationship between the average temperature and average discharge throughout the study period. Subsequently, a minimum positive threshold was set to remove low frequency detections based on the LOD determined in the current study before eDNA concentrations were averaged across the 4 biological and lab replicates from each sampling point. The following calculations were done to calculate the total number of copies per litre (*C*_*L*_):

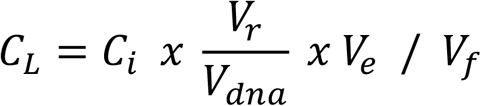

With:

- *C*_*i*_ = Initial concentration on the dPCR
- *V*_*r*_ = The total reaction volume for the dPCR
- *V*_*dna*_ = Volume of DNA in the dPCR
- *V*_*e*_= Volume of DNA extract
- *V*_*f*_= Volume of water filtered

### eDNA downstream transport model

We used a recently developed process-based model (Pont, 2024) that predicts, in a lotic environment, the downstream transport of environmental DNA (eDNA) as a function of downstream retention, degradation processes, and hydraulic conditions. The model assumes that the deposition rate of eDNA on bottom sediments is accurately estimated by the processes governing the sedimentation velocity of fine particles. The predicted eDNA concentration *C*_*d*_(copies /L) at a distance *d* from an eDNA input in the river at a constant concentration of *C*_*0*_ is given by:

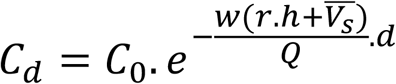

with:

- 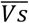 the average eDNA settling velocity (mm.s^-1^)
- *W, h* and *Q* the river width (in m), the river depth (in m) and the discharge ((in m^3^.s^-1^) at a given river section (in m)
- *r*, the rate of eDNA degradation (per hour) which increases exponentially with temperature

The lower and upper limits of the 95% prediction interval around the predicted value are also available. eDNA distance and concentration were computed using the published R script (see (Pont, 2024), Supplement Material 5).

At each eDNA sampling date, the hydraulic parameters (w, h and Q) and water temperature measured at the sites were interpolated every 100 m, to allow the computation of the eDNA concentrations (and their 95% prediction interval), with C_0_ representing the eDNA concentration measured at the source point. When the eDNA concentration at the source point was not at its maximum, the simulation was initialized from the distance to the eDNA source where the concentration was highest (5 dates, between 300 and 600 m from the source point).

The maximum distance is the distance over which the number of eDNA copies per litre is lower than the LOD per litre (LOD_L_) with (Wilcox et al., 2016):

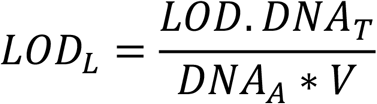

with *V* the mean water volume (in litre) of the eDNA sample, DNA_T_= 100 μL the volume of the DNA elution (μL) produced by extraction and purification, DNA_A_ = 1μL the volume of aliquot of template DNA taken from DNA_A_ for each replicate, and LOD the limit of detection (Pont, 2024).

Since the observed and predicted environmental DNA values were spatially dependent, we tested whether the mean difference between the observed and maximum predicted values (upper limit of the 95% confidence interval) differed significantly from 0. Subsequently, dependent sample sign tests from the “BSDA” package v1.2.2, (Arnholt & Evans, 2017) to assess the fit between the model and the observations. Sign tests were conducted overall and across two distances < 2 km (before the confluence) and subsequently > 2km (after the confluence).

## Results

### Seasonal variation in environmental conditions

Throughout the duration of the sampling period the average water temperatures ranged from 3 ^°^C in January to 15.7 ^°^C in August. Meanwhile the average stream discharge ranged from 0.034 (m^3^/s) in July to 0.926 (m^3^/s) in January (Table S2). Subsequent correlation tests revealed a highly significant negative correlation between discharge and temperature throughout the study period (Spearman’s rank: S = 564, RHO = -0.972, P = <0.001, Figure 2).

**Figure 2:**
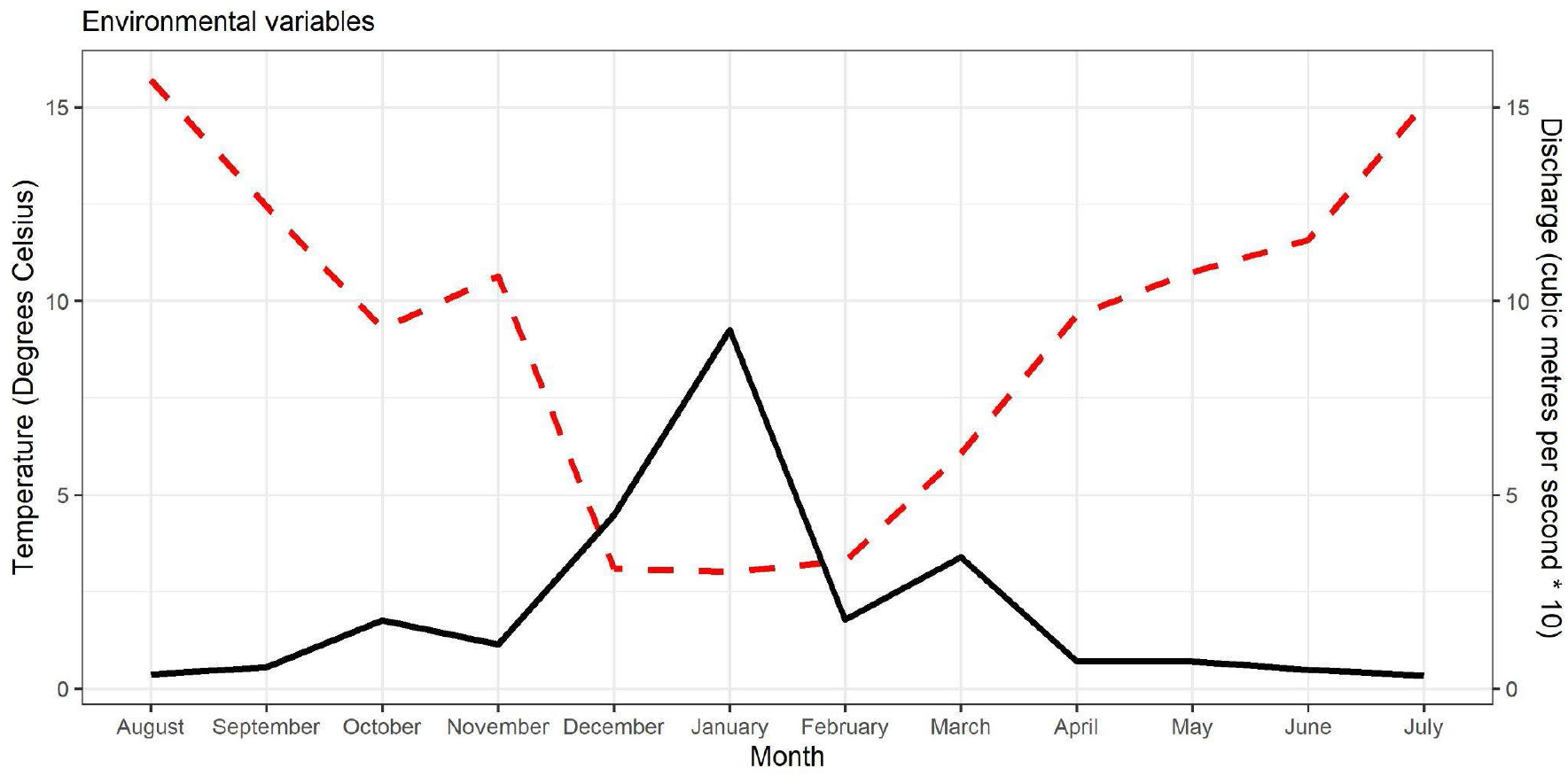
Average discharge (solid black line) and temperature (dashed red line) over the course of the year.

### Limit of Detection

The limit of detection for the current assay was quantified to 1.5 copies per microlitre where 100% (20/20) of reactions successfully amplified (Figure 3). For the tissue concentration at 1.1 copies per microlitre, 90% (18/20) of reactions were successfully amplified. Therefore, a minimum positive threshold of 1.5 copies per microlitre (≧4 partitions) was applied to eDNA samples.

**Figure 3:**
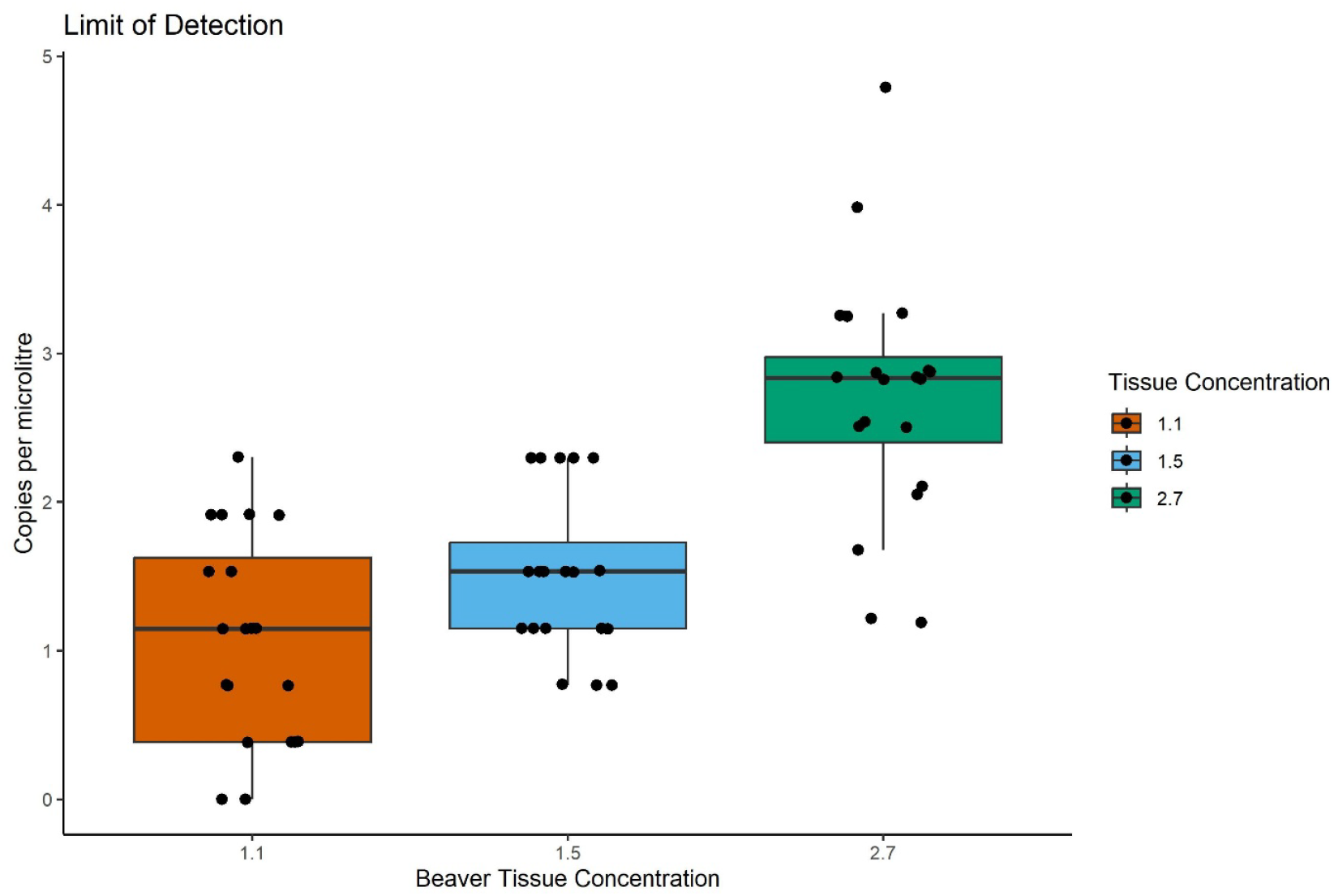
Limit of detection for the Eurasian beaver assay across three different concentrations (n = 20 per tissue concentration).

### Beaver eDNA

No beaver eDNA was detected in any of the 74 controls throughout the 12-month sampling period and no inhibition was observed; the internal positive control amplified successfully in every sample. Furthermore, beaver eDNA was not detected from the control site upstream of the confluence. Beaver eDNA concentration at the source point ranged from 5711.7 - 757.8 copies per litre with the peak in March followed by June. Beaver eDNA was always detected at sites one to four up to 900 metres away throughout all 12 months (Figure 4). The shortest observed eDNA transport was in August and July where beaver eDNA was not detected at site five which was 1.4 km away from the enclosure site. Contrastingly in October, December, January, February, March and May beaver eDNA was consistently detected at the bottom site nine which was 5.8 km downstream of the source point. The shortest predicted eDNA transport was in August where the beaver eDNA signal was below the limit of detection 600 metres from the enclosure (Table S3). The maximum predicted eDNA transport was in January where the eDNA signal was predicted to reach 18.6 km (Table S4), however in October, December, January, February and March beaver eDNA was predicted to reach the Beauly Firth 7.1 km downstream (Table S3).

**Figure 4:**
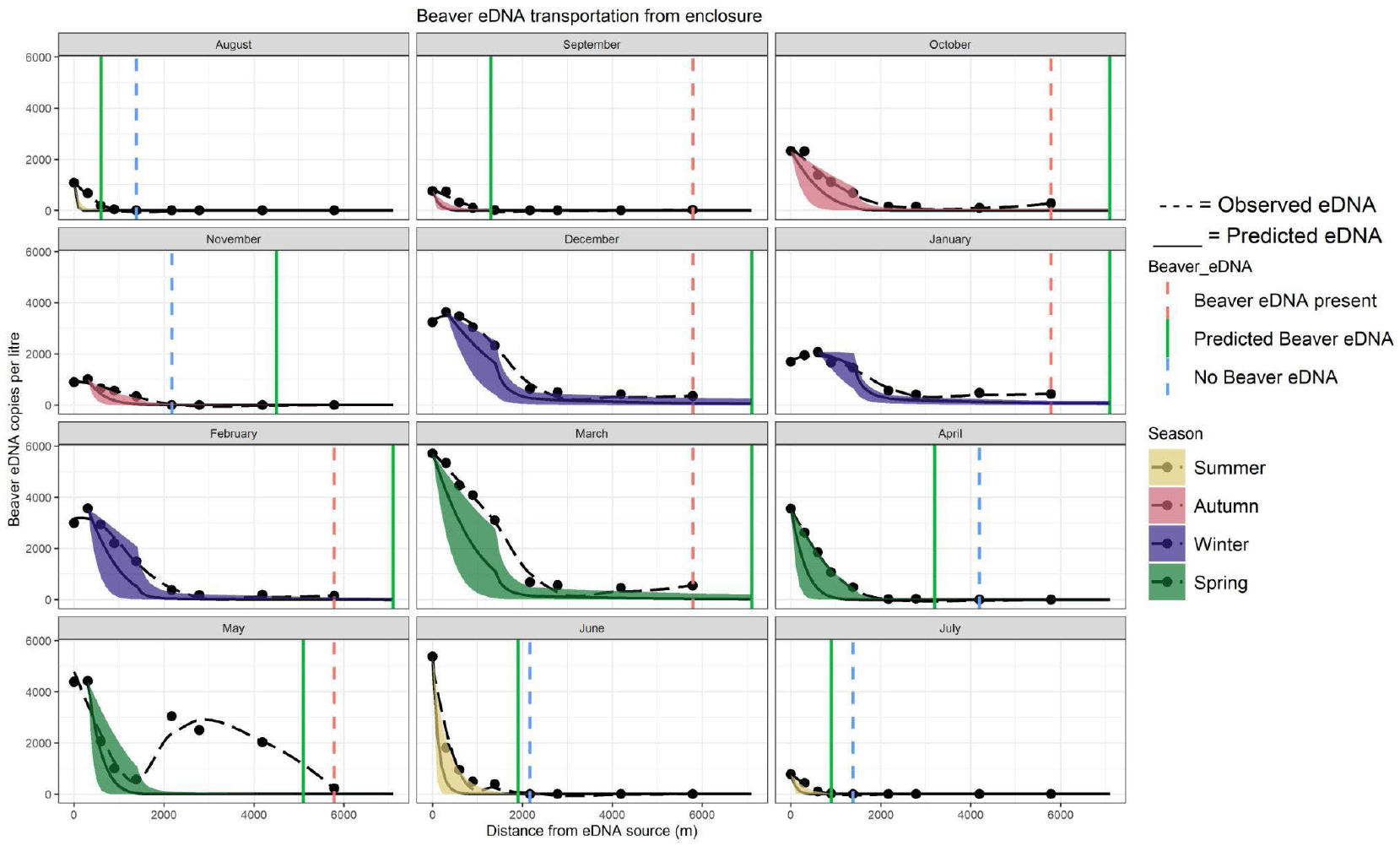
Observed (dashed black line and black dots) and predicted (solid black line) eDNA transport distances each month. The shading represents 95% confidence intervals for the predicted transport distance, and the colour corresponds to the season (yellow: Autumn, red: Spring, purple: Summer, green: Winter). The vertical lines represent the distance beaver eDNA is transported (red: beaver eDNA present, green: maximum predicted beaver eDNA, blue: no beaver eDNA present).

Across the entire river system, the observed eDNA concentrations were significantly higher than the maximum predicted eDNA concentrations (Dependent Sign Test, S = 45, Median = 56.086, P = <0.001). Subsequent examination of the residuals revealed no significant differences between the observed and maximum predicted eDNA concentrations prior to the confluence (< 2km) (Dependent Sign Test, S = 24, Median = 40.493, P = 0.268). However, following the confluence (> 2km) the observed eDNA concentrations were consistently significantly higher than the maximum predicted eDNA concentrations (Dependent Sign Test, S = 21, Median = 69.991, P = <0.001) (Figures S3 + S4).

## Discussion

This study highlights the impact of hydrological and environmental conditions on eDNA transport dynamics. The clear correlation between the predicted and observed eDNA transport provides further evidence towards the utility of eDNA transport models even in smaller river systems and on semi-aquatic mammals. Consistent with our initial prediction; the shortest eDNA transport distances were recorded throughout the summer months, meanwhile the longest eDNA transport distances were seen from December to March where beaver eDNA was predicted to reach the Beauly Firth 7.1 km downstream. Irrespective of environmental conditions, we consistently detected beaver eDNA up to 900 metres downstream of the enclosure site throughout all 12 months of the year. Finally, the observation of beaver eDNA increasing downstream of the enclosure in May highlights the need to consider species-specific behaviours such as dispersal when interpreting the results of these eDNA transport models.

### Seasonal variation in eDNA transport

In the current study, the shortest eDNA transportation (< 1.4 km) was observed during August and July which coincided with the lowest discharge and highest average temperature. This supports the findings from caged brown trout experiments where at low flows the eDNA concentrations observed a sharp decrease with distance from the source point, meanwhile at the highest flows eDNA copies remained consistent throughout the sampled stretch (Jane *et* al., 2015). However, the sampled stretch was only 200 metres (Jane *et al*., 2015), and interestingly our study observed a very similar pattern where the sharpest initial decrease was in June where the eDNA concentration drops by 66% over a 305 metre distance between the source point and site two. Conversely, at higher flows and lower temperatures in October we found a very small decline of 0.8% in the beaver eDNA concentrations over the first 300 metres, and in fact from November to February we observed an increase in the number of eDNA copies between the source point and site two. Our findings of increasing eDNA concentrations a short distance (< 600 m) downstream of the source point at high flows and low temperatures is not unheard of and is discussed in the review conducted by Rourke *et al*., (2022). This initial increase downstream has been described as a breakout phase where the particle mixing and breakdown result in a more homogenous spread of eDNA throughout the water column which leads to an increase in eDNA concentrations (Wood et al., 2020). Previously this breakout phase has been reported over short distances immediately downstream of the source point (< 100 metres) (Bowen et al., 2024; Itakura et al., 2020; Wood et al., 2020). In contrast our study reports a breakout phase of 300 metres for 4 months and at 600 metres in January which coincides with the peak discharge and lowest temperatures. This finding highlights the influence of streamflow and temperature on the breakout phase.

Streamflow and temperature largely influenced the eDNA transport in our study. The longest eDNA transport distances and highest detections of beaver eDNA were positively correlated with discharge and negatively correlated with temperature throughout winter months. This aligns with several previous eDNA metabarcoding studies which documented higher detection probabilities for freshwater fish communities during winter (Griffiths et al., 2025; Milhau et al., 2021). This has primarily been attributed to increased flow and lower temperatures which results in longer eDNA transport and therefore a higher likelihood of detection. Similarly, a comparative analysis between camera trapping and eDNA metabarcoding revealed significantly higher detection efficiency for vertebrates during the high rainfall season (Bai *et* al., 2025). Furthermore, in French Guiana, the higher discharge and lower water temperatures during the rainy season resulted in the spatial homogenisation of beta diversity between sites (Condachou et al., 2025). Subsequent research indicated taxa specific effects of rainfall on vertebrate species richness, where terrestrial species richness significantly increased following rainfall (Reves *et al*., 2026). Meanwhile, aquatic species richness showed signs of decrease however differences were non-significant (Reves et al., 2026). In contrast, single-species assays revealed lower eDNA detection probabilities of an invasive clam (*Corbicula fluminea*) during high streamflows and flood events which resulted in false negatives despite the species being common (Curtis et al., 2021). This study also found a weak positive relationship between eDNA concentration and temperature, however their seasonal analyses revealed that the combined effect of temperature and streamflow had a stronger effect on eDNA concentrations and detectability (Curtis et al., 2021). Interestingly, in our study a dilution effect was observed during the peak flows in January where the maximum beaver eDNA concentration was approximately 42% lower than the concentrations in December and February. However, irrespective of this dilution effect, beaver eDNA could still be detected throughout the entire system up to 5.8 km downstream.

The current study consistently detected beaver eDNA up to 900 m downstream of the enclosure site throughout all 12 months. Similar research from Washington state could reliably detect upstream American beaver presence in 92.4% of eDNA samples taken downstream between 1-3 months post release, the maximum transport distance recorded in this study was 2.9 km however seasonal variation was not considered in this study (Burgher et al., 2024). The peak beaver eDNA concentrations at the source point were observed in March and June with the highest average concentration in Spring, which likely coincides with beavers increasing their activity following the colder months (Korbelová et al., 2016; Mott et al., 2011). The large eDNA concentrations in June could represent an increase in eDNA following the birth of kits, which research from Norway has previously been shown to peak in late May (Mayer et al., 2017b). Similar prior eDNA surveys have demonstrated a peak in the airborne eDNA signal for pocket gophers (*Geomyidae*) during the breeding season (Johnson et al., 2023). The increase in beaver eDNA < 2 km downstream of the enclosure site during May could represent an escapee from the enclosure site. This would coincide with the dispersal window for juvenile Eurasian beavers (> 20 months) from their natal range (Campbell-Palmer et al., 2021a; Hartman, 1997; Rosell et al., 1998), or it could represent explorative trips by adult beavers outside the enclosure (Mayer et al., 2017a). Escapes from captive beaver populations have been reported throughout the UK (Campbell et al., 2012) and mainland Europe (Halley et al., 2021). Nonetheless, there was no field sign evidence or eDNA signal in subsequent months to suggest this potential escapee had remained in the system. Therefore, future eDNA surveys to study beaver distribution should consider avoiding sampling during spring to mitigate the signal from dispersing individuals and ensure the most accurate distribution of beavers.

### Performance of the eDNA transport model

Throughout the study, the primary model parameters which varied temporally were the average discharge and average temperature which experienced a 27-fold increase and 5-fold reduction respectively between July and January (Table S2). Nonetheless, with the exception of May, the predicted eDNA transport model reliably reproduces the decrease in eDNA concentrations downstream of the source point over the course of the year. There were however challenges following the confluence between the small river and Moniack river. The initial dilution in eDNA concentrations is well predicted by the eDNA transport model, however, the model underestimates the eDNA concentrations over the subsequent 3.6 km particularly when the discharge is high (Figures S3 + S4). In the Danube river, predicted concentrations of brown trout (*Salmo trutta*) were slightly lower than observed eDNA concentrations, which was attributed to additional small inputs of eDNA from tributaries further downstream which could not be accounted for (Pont, 2024). In the current study, no wild beaver population is known downstream of the enclosure and no beaver eDNA was detected from the control site upstream of the confluence. Therefore, additional downstream inputs of beaver eDNA are unlikely, however they cannot be ruled out which might explain the disparity between the predicted and observed eDNA concentrations.

Another explanation behind the disparity in the model accuracy could be a change in decay rate between the small river and the larger Moniack river (Barnes et al., 2014). The small river is a slow-flowing agricultural river with an average width and depth of 2.3 metres and 13 cm respectively; the substrate primarily consists of small pebbles and fine sediment (Figure S5). In contrast the Moniack river is faster flowing with an average width and depth of 5.1 metres and 16 cm respectively and the substrate is predominantly cobbles (Figure S5). Rapid transport of eDNA has been highlighted in cobble sections where no decrease in eDNA concentrations were recorded, contrastingly the shortest transport distances were observed in fine sediment sections likely due to increased instream retention (Shogren et al., 2017). In our study, the smaller substrate in the small river results in high eDNA deposition and instream retention causing a shorter eDNA transport distance. This was particularly clear in the summer where beaver eDNA could not be detected 2.1 km downstream at site six. Conversely, the larger cobble substrate in the Moniack river results in minimal eDNA deposition and subsequent rapid flushing which would explain the larger than predicted transport distances observed in this study (Jerde et al., 2016; Shogren et al., 2017) (Figure S2 + Figure S5). This highlights the challenges in accounting for changes in hydrological and substrate characteristics surrounding confluences to ensure an accurate prediction of eDNA transport.

## Conclusions

Our findings reveal the influence of seasonal environmental conditions on eDNA transport dynamics and highlight the importance in accounting for this when designing future studies and monitoring programmes. To study the localised distribution of species, we recommend sampling in the summer during low flow conditions and high temperatures when eDNA transport away from the source is minimal. In the context of monitoring beaver distribution, this approach would help to narrow down the sections of a catchment where beavers are most active. Conversely to maximise species detections, we suggest sampling in the winter which had the longest eDNA transport distances and highest detectability over a long range of the catchment. This approach is particularly suitable when the objective of monitoring is to determine whether beavers have successfully colonised a catchment or tributary. Our results in May highlight the need to consider species-specific behaviours such as dispersal into eDNA studies. Finally, our results support the integration of eDNA transport models in smaller river systems whilst reinforcing the need to consider stream specific decay rates surrounding confluences.

## Supporting information

Word document containing all supporting information for the publication

## DATA AVAILABILITY STATEMENT

All scripts and corresponding data have been archived and made available at Zenodo: https://doi.org/10.5281/zenodo.18485541

