## Supplementary material for "Process-based model predicts seasonal variation in eDNA transport - a case study on Eurasian beavers in a small river": Word document containing all supporting information for the publication

**Methods S1:** Detailed lab workflow

**Sample collection:**

Between 06/08/2024 - 03/07/2025, 218 2-litre surface samples were taken using sterile Gosselin HDPE plastic bottles (Fisher Scientific UK Ltd., Loughborough, UK). A field blank consisting of 2-L purified water was used each day (n=13), this was taken into the field and handled alongside eDNA samples to monitor for contamination. Following collection samples were placed on ice packs within bleach sterilised coolboxes before being taken back to an eDNA filtration facility at the University of Highlands and Islands.

**Filtration**

Water samples were vacuum filtered within 24 hours of collection following the methodology described in Griffiths *et al*., (2023). Briefly, 2-L samples were vacuum filtered through sterile 0.45 μm mixed cellulose nitrate membrane filters with pads (47 mm diameter; Whatman, GE Healthcare) using Pall filtration units. Two filters were used for each sample to minimise clogging, and up to 30 minutes per filter was allowed for 1-L of water to pass through. Filters were removed from pads using sterile tweezers, rolled, and placed back-to-back in sterile 5 ml Axygen screw-cap tubes (Fisher Scientific UK Ltd., Loughborough, UK) before being stored at -20 °C. Up to nine samples were filtered each run and blanks (n=13) were filtered during the last round of each day. Between runs, attachments were submerged in 10% v/v chlorine-based bleach solution (Fisher Scientific UK Ltd., Loughborough, UK) for 10 min, soaked in 5% v/v MicroSol detergent (Scientific Laboratory Supplies, Newhaven, UK) for a further 5 min, before being rinsed carefully with purified water to remove any remaining detergent. Throughout filtration, all benchtops and equipment within the filtration lab were sterilised using 10% bleach solution and cleaned using 70% v/v ethanol (Fisher Scientific UK Ltd., Loughborough, UK) (Griffiths *et al*., 2023).

**DNA extractions**

DNA extractions were carried out following the mu-DNA lysis water protocol described in Sellers *et al*., (2018) but with the following modifications: instead of garnet beads, the lysis stage used an incubation based method where 34 μl of 20 mg/mL proteinase K (VWR, Leicestershire, UK) was added to 750 μl of lysis solution and 250 μl of water lysis additive. The filter was then pressed to the bottom of the tube using the pipette tip until it was submerged below the liquid before tubes were incubated at 55°C overnight. The ethanol wash step was repeated twice to minimise the chances of inhibition before the rest of the protocol was followed as normal (Sellers *et al.*, 2018). An extraction blank (n=12), consisting only of extraction buffers, was extracted alongside samples to monitor for contamination. Following extraction, the eluted DNA extracts (100 µL) were visualised using the QIAxpert (Qiagen, Hilden, Germany) to confirm DNA was successfully isolated, then stored at −20°C.

**Digital PCR**

Digital PCR was performed in duplicate (2 x PCR replicates per eDNA extract) using the following unpublished primers F (5'-CCTCGGTGCCATCAACTTTA-3') and R (5'-GCAGTAACTAGGACGGATCATAC-3') to amplify a 102 bp fragment of the Eurasian beaver COX1 gene (Bouhouche, 2023). These primers were previously validated *in vitro, in silico* and *in situ* to ensure high specificity to the Eurasian beaver using qPCR (Bouhouche, 2023). A positive control (quantified at 0.01 ng/µl) of Eurasian beaver tissue and a negative control of molecular grade water were used for each plate and the TaqMan Exogenous Internal Positive Control was used to test for inhibition (Applied Biosystems, Warrington, UK). Reaction volumes consisted of 13.3 µl QIAcuity EG PCR kit (QIAGEN, Manchester, UK) 1 µl primers (0.5 x each 10 µM Primer (Integrated DNA Technologies, Belgium), 4 µl IPC Mix and 0.4 µl IPC DNA (Applied Biosystems, Warrington, UK), 16.3 µl of molecular grade water and 5 µl of DNA extract.

Reactions were run on the QIAcuity-00896 digital PCR system (QIAGEN, Manchester, UK) using the 26k nanowell plates with the following thermocycling profile: 95°C for 10 min, 35 cycles of 95°C for 15 s and 60°C for 1 min. The beaver and IPC assays used the EVAGreen dye and VIC dye respectively and imaging for both channels was conducted at an exposure duration of 300 ms and gain 6. As the EVAGreen assay fluoresces both beaver eDNA and the IPC a common threshold was set at 130 RFU for the beaver channel to ensure only beaver eDNA was retained (Figure S1), meanwhile the auto-threshold was used to quantify the IPC. The limit of detection (LOD) was calculated following a discrete threshold approach which ran three different concentrations of beaver tissue aiming to achieve ≧ 95% detections across 20 replicates (Klymus *et al.*, 2020).

### Methods References

Bouhouche, M. (2023). Environmental DNA analyses for detecting the Eurasian beaver (*Castor fiber*) and assessing biodiversity: a metabarcoding and qPCR study (MSc). University of South-Eastern Norway Faculty of Technology, Natural Science and Maritime Science, Kongsberg, Norway.

Griffiths, N. P., Wright, R. M., Hänfling, B., Bolland, J. D., Drakou, K., Sellers, G. S., … Vasquez, M. I. (2023). Integrating environmental DNA monitoring to inform eel (Anguilla anguilla) status in freshwaters at their easternmost range-A case study in Cyprus. *Ecology and evolution*, *13*, e9800.

Klymus, K. E., Merkes, C. M., Allison, M. J., Goldberg, C. S., Helbing, C. C., Hunter, M. E., … Richter, C. A. (2020). Reporting the limits of detection and quantification for environmental DNA assays. *Environmental DNA*, *2*, 271–282.

Sellers, G. S., Di Muri, C., Gómez, A., & Hänfling, B. (2018). Mu-DNA: a modular universal DNA extraction method adaptable for a wide range of sample types. *Metabarcoding and metagenomics*, *2*, e24556.

**Table S1:** Elevation and distance from coast for the 10 sites sampled throughout the study

| Site | Elevation (m) | Distance from Source (M) |
|---|---|---|
| S1 | 155 | 0 |
| S2 | 145 | 305 |
| S3 | 145 | 597 |
| S4 | 141 | 896 |
| S5 | 137 | 1385 |
| S6 | 82 | 2171 |
| S7 | 48 | 2785 |
| S8 | 18 | 4193 |
| S9 | 4 | 5789 |
| C | 131 | n/a |

**Table S2:** Average discharge ($m^3/s$) and temperature (℃) measured across the 12 months

| Month | Discharge ($m^3/s$) | Temperature (℃) | Order visited | Season | Date |
|---|---|---|---|---|---|
| August | 0.036 | 15.71 | 1 | Summer | 06/08/2024 |
| September | 0.056 | 12.44 | 2 | Autumn | 09/09/2024 |
| October | 0.177 | 9.28 | 3 | Autumn | 09/10/2024 |
| November | 0.115 | 10.64 | 4 | Autumn | 07/11/2024 |

| | | | | | |
|---|---|---|---|---|---|
| December | 0.448 | 3.10 | 5 | Winter | 03/12/2024 |
| January | 0.926 | 3.01 | 6 | Winter | 03/01/2025 |
| February | 0.178 | 3.28 | 7 | Winter | 12/02/2025 |
| March | 0.339 | 6.07 | 8 | Spring | 11/03/2025 |
| April | 0.071 | 9.63 | 9 | Spring | 10/04/2025 |
| May | 0.071 | 10.76 | 10 | Spring | 06/05/2025 |
| June | 0.048 | 11.58 | 11 | Summer | 05/06/2025 |
| July | 0.034 | 15.07 | 12 | Summer | 03/07/2025 |

**Table S3:** Observed and predicted eDNA transport distances in KM, a distance of 7.1 km represents beaver eDNA reaching the Beauly Firth

| Month | Observed eDNA distance (km) | Minimum predicted eDNA distance (km) | Mean predicted eDNA distance (km) | Maximum predicted eDNA distance (km) |
|---|---|---|---|---|
| August | 0.9 | 0 | 0.2 | 0.6 |
| September | 5.8 | 0.1 | 0.4 | 1.3 |
| October | 5.8 | 0.8 | 1.8 | 7.1 |
| November | 1.4 | 0.7 | 1.5 | 4.5 |
| December | 5.8 | 2.6 | 7.1 | 7.1 |
| January | 5.8 | 4.1 | 7.1 | 7.1 |
| February | 5.8 | 1.2 | 3.4 | 7.1 |
| March | 5.8 | 1.5 | 7.1 | 7.1 |
| April | 2.8 | 0.4 | 1.2 | 3.2 |
| May | 5.8 | 0.7 | 1.4 | 5.1 |
| June | 1.4 | 0.2 | 0.7 | 1.9 |
| July | 0.9 | 0.1 | 0.3 | 0.9 |

**Table S4:** Maximum eDNA transport distance calculated from the peak eDNA concentrations each month and from a single set of environmental conditions.

| **Month** | **Predicted eDNA distance (km) from the source point** |
|---|---|
| August | 0.3 |
| September | 0.9 |
| October | 5.7 |
| November | 2.9 |
| December | 16.9 |
| January | 18.6 |
| February | 5.8 |
| March | 11.9 |
| April | 2.8 |
| May | 2.9 |
| June | 1.1 |
| July | 0.7 |

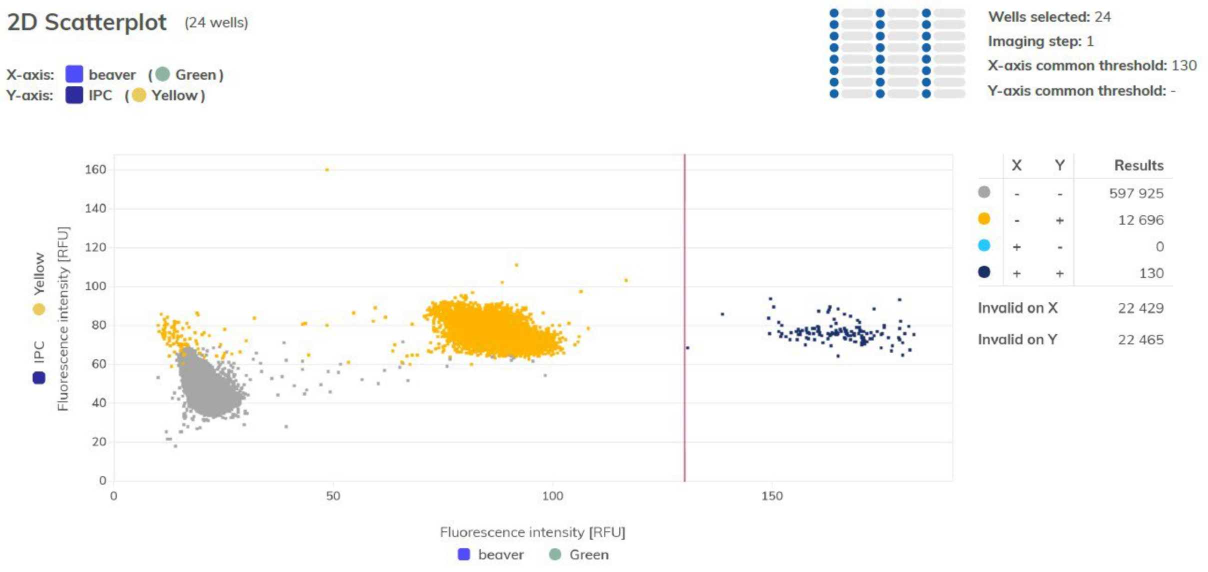


**Figure S1:** 2D-Scatterplot generated via QIAcuity software suite 3.2.0.0, displaying the relative fluorescence intensity from each marker. The red line highlights the common threshold set at 130 to determine positive beaver amplification.

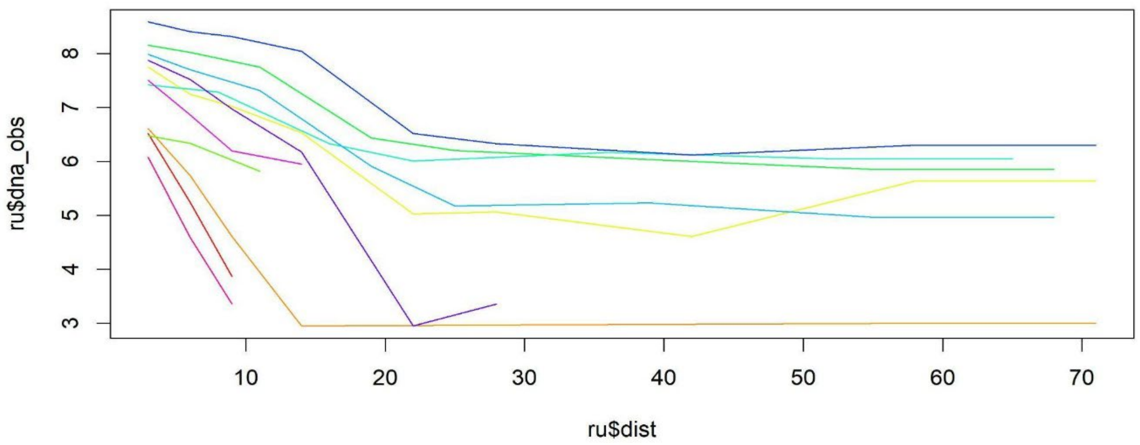


**Figure S2:** Log transformed relationship between the observed eDNA concentrations and distance from source point. May has been omitted from this plot due to the increase in beaver eDNA downstream of the enclosure site.

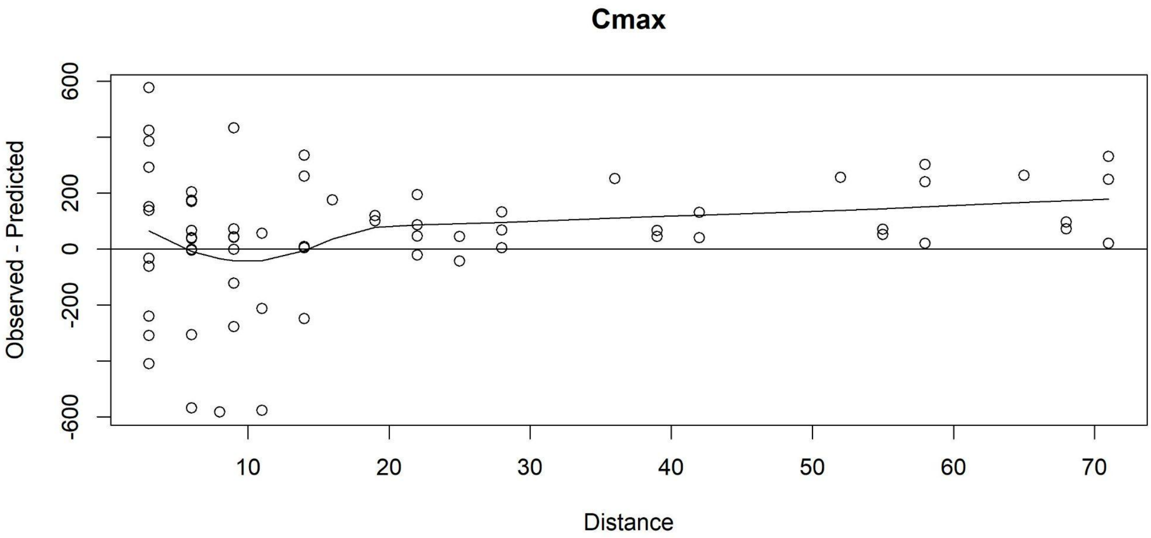


**Figure S3:** Residual plot of the observed eDNA - predicted eDNA concentrations against distance.

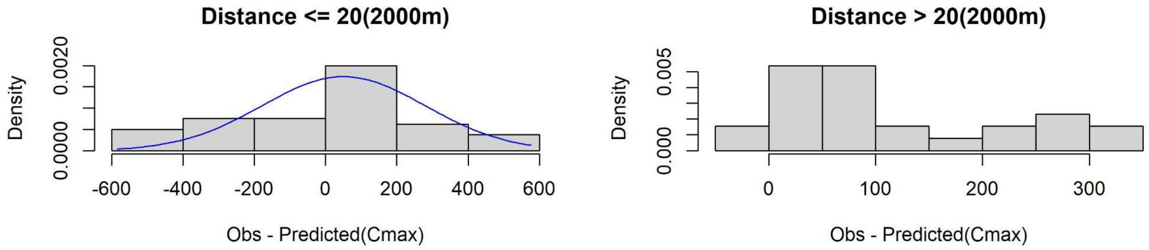


**Figure S4:** Comparative histograms displaying the difference in the observed and predicted eDNA concentrations before (< 2 km) and after (> 2 km) the confluence.

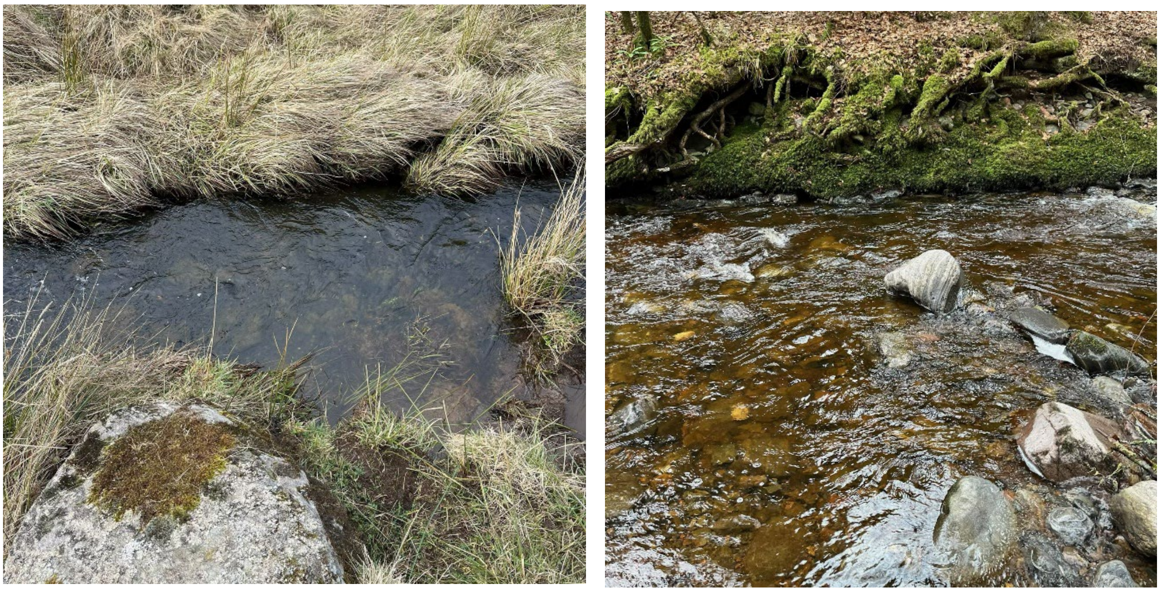

**Figure S5:** Comparative pictures from March 2025 displaying the difference in hydrological and substrate conditions between the small river (left) and Moniack river (right)
